# Calsequestrin localization at RyR2 clusters enables calcium wave propagation in ventricular myocytes

**DOI:** 10.64898/2026.06.15.732280

**Authors:** David Conesa, Blas Echebarria, Leif Hove-Madsen, Yohannes Shiferaw, Enrique Alvarez-Lacalle

## Abstract

Intracellular calcium waves in cardiac myocytes propagate through a fire-diffuse-fire mechanism in which calcium released from one RyR2 cluster diffuses to neighboring clusters and triggers their activation. Yet propagation faces a fundamental physical difficulty: the calcium signal must cross distances of 1–2 µm between Z-planes while being attenuated by cytosolic buffering and diffusion, and at the same time the release site depletes its local sarcoplasmic reticulum calcium store. How waves propagate efficiently despite these constraints has remained unclear. We developed a three-dimensional computational model of mouse ventricular myocytes at 100 nm resolution to address this question. Our central finding is that co-localization of calsequestrin2 (CASQ2) with RyR2 clusters is required for robust wave propagation. In a physiological model, where CASQ2 is concentrated at release sites as observed experimentally, calcium waves propagate reliably across the cell with velocities that match the experimental range. In contrast, when CASQ2 is distributed uniformly throughout the sarcoplasmic reticulum, keeping total CASQ2 unchanged, the wavefront stalls. These results identify CASQ2–RyR2 co-localization as a key structural requirement for effective calcium wave propagation in ventricular myocytes.

**Author summary:** Calcium waves in cardiomyocytes are thought to underlie the onset of malignant cardiac arrhythmias, such as ventricular tachycardia and fibrillation. Yet, the specific conditions that regulate the transition from local calcium sparks to sustained waves remain poorly understood. Using a newly developed computational model of calcium handling, we demonstrate that the spatial distribution of key regulatory proteins is a critical determinant of arrhythmogenicity. Specifically, we found that calsequestrin2, which buffers Ca2+ within the sarcoplasmic reticulum, must be strictly colocalized with Ca2+ release proteins to facilitate sustained wave propagation. This discovery suggests that cardiac stability depends less on the total quantity of protein and more on its precise architectural organization. The consequences of this finding are significant: it implies that “spatial dysregulation”—where proteins are present but mislocalized—may be a hidden driver of arrhythmias even when protein levels appear normal. This shifts the therapeutic focus from simply altering ion channel conductance to preserving or restoring the structural tethering of the junctional SR. By focusing on the nanodomain architecture, we can better understand how cellular remodeling leads to life-threatening electrical instability.

## Introduction

Cardiac myocytes generate spontaneous intracellular calcium waves under conditions of calcium overload [1–4]. The likelihood of a spontaneous release is strongly dependent on the luminal calcium concentration within the SR, with higher SR calcium load increasing the probability of waves. The sensitivity of the ryanodine receptor (RyR2) to luminal calcium can affect this threshold for spontaneous waves [5, 6], but the general structure of the propagation is the same. In these events, calcium released from the sarcoplasmic reticulum at one RyR2 cluster diffuses to neighboring release sites and triggers further release, producing a regenerative wave that travels through the cell [7, 8]. These waves have a dual physiological significance [9, 10]. On one hand, they can act as a homeostatic mechanism that helps clear excess calcium by promoting extrusion through the sodium-calcium exchanger (NCX) [11]. On the other hand, that same extrusion generates inward current, which can depolarize the membrane and promote delayed afterdepolarizations and triggered activity [12, 13]. Understanding how calcium waves propagate is therefore essential for linking subcellular calcium dynamics to cellular electrical stability.

The underlying mechanism of propagation is known as the fire-diffuse-fire mechanism [14] where calcium released by one RyR2 cluster raises local cytosolic calcium that then diffuses to neighboring sites and, if the calcium signal is large enough, triggers opening of these sites and release of more calcium [15]. Although this mechanism is conceptually simple, its physical realization in a cardiac cell is difficult and complex. Cytosolic buffers rapidly bind free calcium and reduce the amplitude of the mobile calcium signal [16]. Moreover, diffusion spreads calcium in three dimensions and therefore dilutes the signal as it moves away from the source [17]. In addition, the spacing between successive Z-planes is approximately 1.6 µm, so the calcium signal must remain sufficiently strong over a substantial distance. At the same time, the active release site depletes the local sarcoplasmic reticulum calcium store, reducing both the luminal free calcium concentration and the release flux through RyR2 [18, 19].

These constraints make wave propagation fundamentally more demanding than local spark generation, which only requires calcium release from a single RyR2 cluster sufficient to increase calcium in the immediate vicinity of the cluster [7]. In contrast, a propagating wave requires each active site to generate a cytosolic calcium signal strong enough to bridge the gap to the next Z-plane and trigger a new release event [20, 21]. This creates opposing requirements because the release must be large and sustained enough to overcome cytosolic attenuation, but also terminate appropriately to avoid persistent or spatially uncontrolled activation. Previous computational models have captured important aspects of calcium sparks and waves [13, 21–27], but many rely on compartmental descriptions that impose effective coupling between release sites rather than resolving the full spatial transport problem directly. Such models can reproduce wavelike behavior, but they do not fully explain how a wavefront survives the combined effects of buffering, diffusion, and local source depletion.

In this work, we developed a three-dimensional model of a mouse ventricular myocyte at 100 nm resolution. This spatially resolved framework allows us to determine directly whether calcium released from one cluster can generate a sufficiently strong signal to activate neighboring clusters across the physical distances and buffering that separate them. Our main finding is that co-localization of calsequestrin (CASQ2) with RyR2 is required for robust wave propagation. CASQ2 is the principal luminal calcium buffer of the sarcoplasmic reticulum [28], and experimental observations indicate that it is anchored near RyR2 clusters by proteins such as triadin and junctin [29,30]. Our simulations show that this spatial organization is functionally essential. Thus, waves propagate robustly when CASQ2 is concentrated at RyR2 release sites, whereas redistributing the same total amount of CASQ2 uniformly throughout the sarcoplasmic reticulum causes the wavefront to stall.

## Materials and Methods

### Experimental data

#### RyR2-calsequestrin localization

Ventricular myocytes were isolated from mice with GFP-tagged RyR2 as previously described [31], allowing for direct visualization of RyR2 clusters in the myocytes using confocal imaging [31]. To visualize the distribution of CASQ2 relative to the RyR2 clusters, isolated myocytes were fixed with 5 % paraformaldehyde for 10 minutes, incubated with PBS / Glycine 0.1 M for 10 minutes and then with PBS / 0.2 % Triton X-100 for at least 15 minutes to permeabilize the cells. Cells were incubated with PBS / 0.2 % Tween 20, and 10 % Horse serum, for 30 minutes to block non-specific sites. CASQ2 was labeled with primary rabbit anti-CASQ2 (1:500, Ab-3516, Abcam). The secondary antibody AlexaFluor 594 anti-rabbit (diluted 1:1000) was used to stain CASQ2 in red. The distributions of RyR2 and CASQ2 were then visualized using sequential confocal imaging. GFP-tagged RyR2 was excited at 488 nm and fluorescence emission collected between 500 and 570 nm. To visualize CASQ2, AlexaFluor 594 was excited at 543 nm and fluorescence emission was collected between 570 and 700 nm. Representative images of GFP-tagged RyR2, immunofluorescently labelled CASQ2 and overlay of the two images are shown in figure 1A.

**Fig 1.**
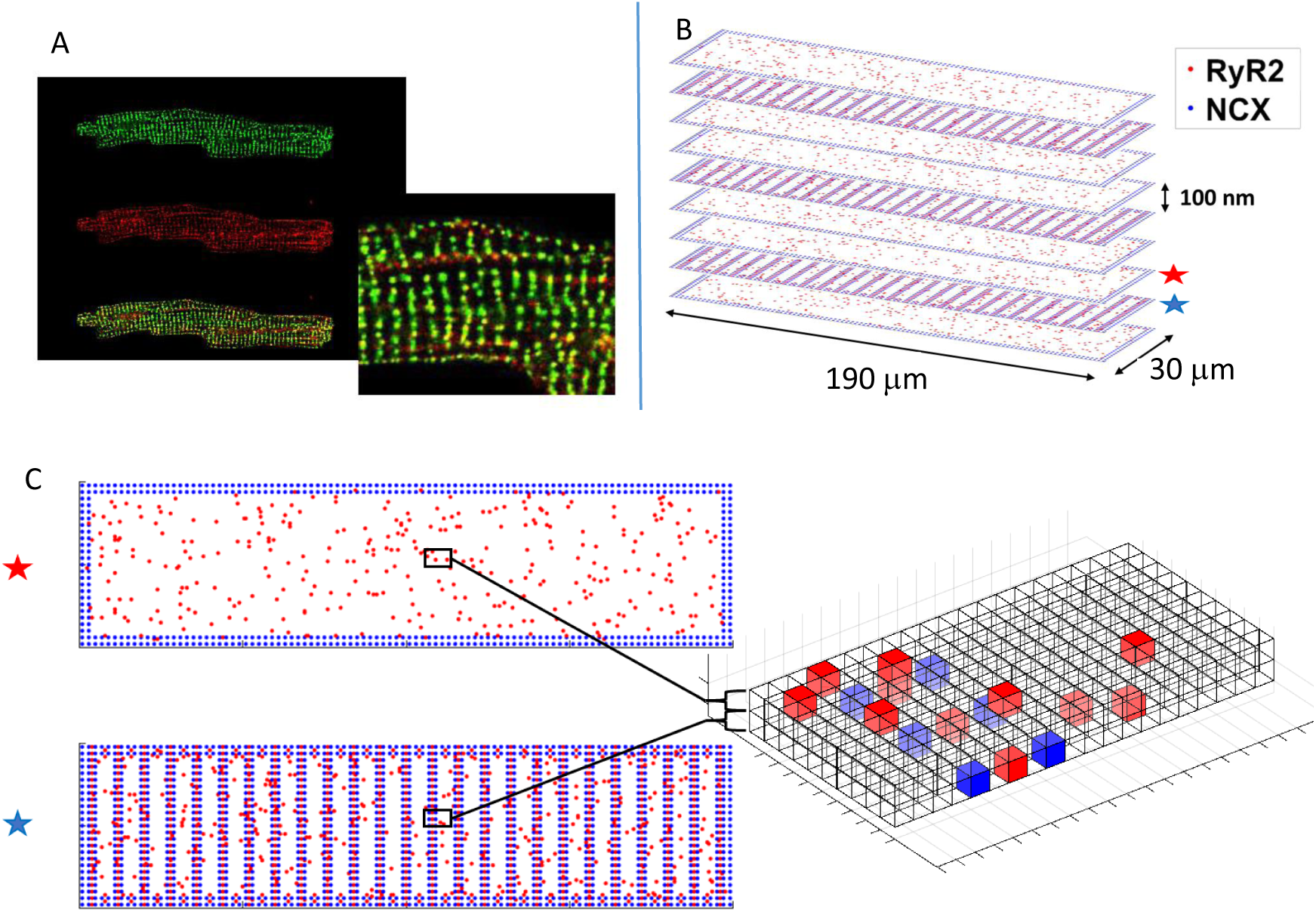
Subcellular 3D structure. Experiments and model. A) Immunofluorescent labeling of RyR2 and CASQ2 showing the distribution of RyR2 (green) and CASQ2 (red) in human isolated atrial cells. B) The full structure of our model cell as 2-D planes stacked to form a 3-D structure showing RyR2 (red) and NCX (blue) distributions. C) Below are two contiguous planes and a 3-D voxel representation of a small region of them.

#### Calcium waves in isolated ventricular mouse

To visualize calcium waves and their propagation in isolated mouse ventricular myocytes, cells were loaded with the fluorescent calcium indicator Rhod-2 (2.5 µM) for 20 minutes followed by wash for at least 30 mins. Rhod-2 was excited at 543 nM with laser power set to 20 % and attenuation to 10 %. Fluorescence emission was collected between 550 and 750 nm. Images (512x64 pixels) were recorded with a resonance scanning confocal microscope (Leica SP5 TCS AOBS) at a frame rate of approximately 80 Hz. Images were analyzed manually using Leica LASAF software or a custummade program (10.1093/cvr/cvv046) to detect calcium waves and determine their propagation velocity and properties.

### Subcellular calcium cycling model

#### Generalities

We employ a deterministic model for calcium cycling that omits compartments and utilizes a higher spatial discretization than typical models based in compartments. This offers greater flexibility in positioning the RyR2 clusters within the z-planes or off-planes [21, 26, 27]. We study an isolated cell kept at resting potential, where LCC flux is negligible but NCX extrusion is not. The model implements RyR2 opening regulation by luminal calcium levels [32, 33], while RyR2 inactivation is influenced by luminal calcium and calcium bound to calmodulin (CaM) [34]. In the following, we present the main features of the model and simulations. Further details of the model and the numerical implementation can be found in the S1 File.

#### Spatial structure of the calcium cycling model

The cell is represented as a 3D grid of voxels, each measuring 100 nm × 100 nm × 100 nm (see Figure 1). A typical full cell consists of 300 × 1900 × 150 voxels. To speed up simulations of all scenarios, we perform most of our simulations in the central layer of a small cell comprising 300 × 1900 × 9 voxels, and use the large cell only to verify the robustness of the main results.

Each voxel contains two distinct domains: the cytosolic and the sarcoplasmic reticulum (SR). Calcium cycling occurs within and between these two domains via the release of calcium from the SR through the RyR2 and uptake by SERCA, with calcium ions diffusing between neighboring voxels in both the SR and the cytosol. Calcium is also extruded via NCX. We define, in each voxel, the total cytosolic calcium concentration c*_T_* as the sum of the free calcium concentration c*_i_* and the calcium bound to cytosolic buffers c*_b_*, i.e., c*_T_* = c*_i_* + c*_b_*. Similarly, we define the total calcium concentration *c*_SR_^TOT^ = c_SR_ + c_SRb_ , which includes the free SR calcium concentration c_SR_ and calcium bound to calsequestrin c_SR_*_b_* . Then, the total calcium concentration in the cell is C*_T OT_* = c*_T_* f*_i_* + *c*_SR_^TOT^ *f_SR_*, with *f_i_* and *f_SR_* the cell fractions of cytosolic and SR volumes.

The basic calcium cycling in each voxel thus reads:

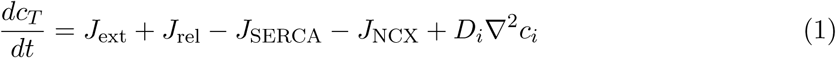

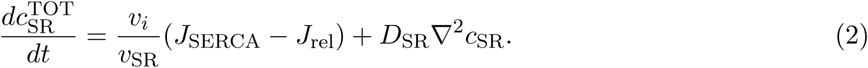

with *v_i_* and *v_SR_* as the volume fractions of the cytosol and SR within the voxel, and *D_i_* and *D_SR_* the diffusion coefficients for calcium in the cytosol and SR, respectively.

#### RyR2 structure and spatial distribution

The distribution of RyR2 along the cell boundary and within the transverse tubular system (T-tubules), facilitates rapid transmission of electrical signals and efficient calcium signaling deep within the cell [35]. T-tubules enter the cell along specific planes known as Z-planes, which are regularly spaced every 1.6 µm along the longitudinal axis of the cell. In our model, every 16nth plane along the longitudinal direction is designated as a Z-plane. Within each Z-plane, clusters of RyR2 are positioned regularly at their average physiological distances of 300-500 nm. Specifically, RyR2 clusters are spaced 400 nm apart in the long transverse direction and 300 nm apart in the short transverse direction (see Fig. 1). Additionally, to account for RyR2 clusters outside the Z-planes, we include an off-plane population of RyR2 clusters by populating 1% of the voxels outside the Z-planes with RyR2. The location of the RyR2 clusters is fixed randomly.

In ventricular cardiomyocytes NCX is located in the Z-planes along the t-tubules and concentrated near the boundaries [36–39]. We position NCX channels within every other voxel located between 200 nm and 800 nm from each border of the cell. Figure 1 illustrates the relative distribution of RyR2 clusters and NCX channels in our model.

In those voxels with RyR, the flux of Ca through the RyR2 is given by

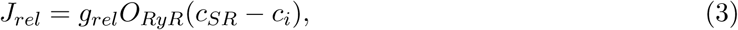

where O*_RyR_*is the fraction of RyR2 in the clusters that are in the open state. Figure 2A shows the cartoon image of two of the four units of a RyR2 channel in the open state and the direction of the release through the pore. We consider a RyR2 model of three states (Fig. 2B), with closed (C), inactivated (I), and open state (O*_RyR_*), which captures the main effects of RyR2 regulation. Defining R = C + O*_RyR_*, the fraction of inactivated states is then 1 −R. Transition dynamics reads:

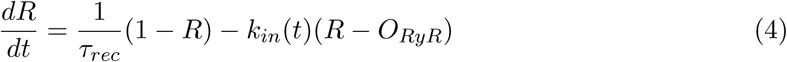

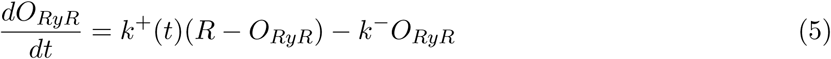

**Fig 2.**
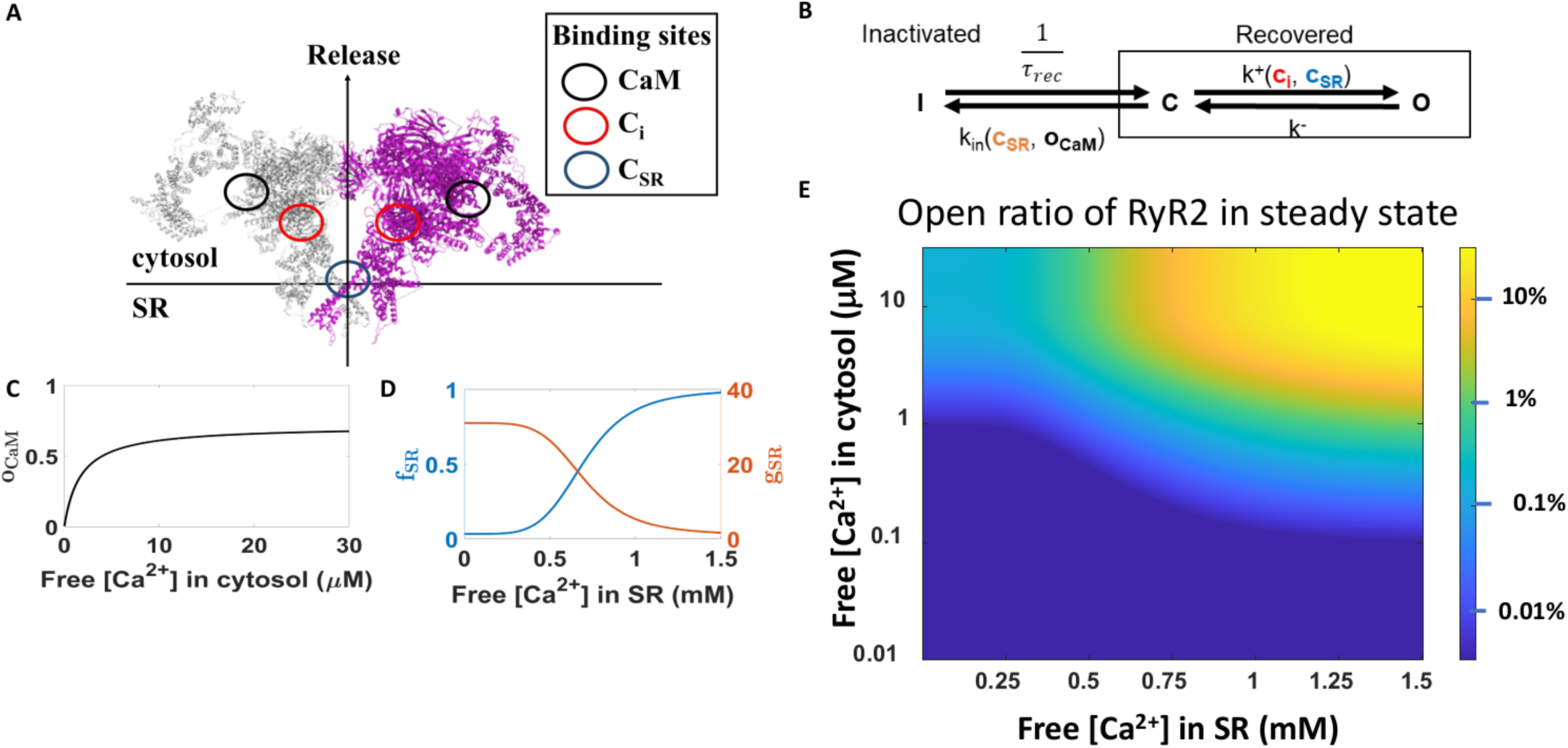
A): 3D structure of human RyR2 (PDB ID: 7UA9 [40–43]). Only two subunits of the tetramer are shown for clarity, indicating the location of the binding sites for calmodulin (CaM), and cytosolic (c*_i_*) and luminal (c*_SR_*) calcium. B): Three-state Markov chain for RyR2 gating, with inactivated (I) and two recovered states, close (C) and open (O). C): Occupation of RyR2 - CaM binding sites in steady state as a function of free cytosolic Ca^2+^. D): Modulatory functions f*_SR_* and g*_SR_* of opening and inactivation of RyR2 by effects of luminal Ca^2+^ binding in steady state as a function of free SR Ca^2+^. E): Logarithmic color-code of the RyR2 open fraction in steady state, at different free cytosolic and SR Ca^2+^ concentrations. The opening rate k^+^ and the inactivation rate k*_in_* are taken at steady state as well.

Close and recovery rates are considered constant values, while open and inactivation rates (k^+^, k*_in_*) are not constant and play a crucial role in the regulation of the RyR2.

#### RyR2 regulation

The opening rate k_+_ of the RyR2 is regulated by cytosolic free Ca^2+^ concentration through cooperative binding of c*_i_*, and luminal calcium concentration c_SR_ [44, 45], through a phenomenological gating variable o_LCA_:

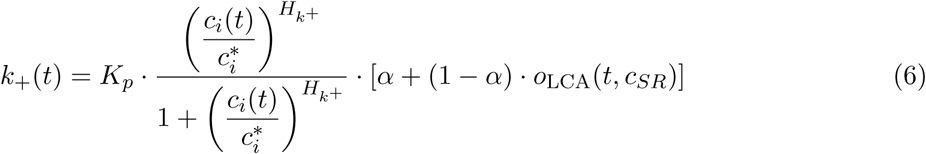

Here K*_p_* is the maximum possible opening rate of RyR2 channels, c^∗^ is the half-saturation constant for cytosolic Ca^2+^, and H*_k_*^+^ is the Hill coefficient representing the degree of cooperativity in cytosolic Ca^2+^ binding. The gate o_LCA_ relaxes, as

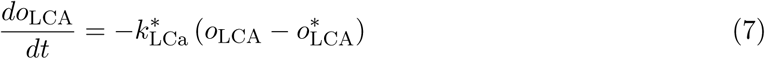

to a steady-state value [46] given by 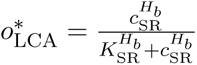, in a time scale that also depends on SR Ca, by 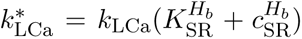. As a result, when luminal calcium decreases, occupancy decreases and the opening rate k_+_ decreases.

The inactivation rate, k_in_, depends on two main factors: luminal calcium via a gate o_LCA_ and the occupancy of the Ca^2+^-CaM complex binding site o_CaM_ [34, 47, 48]:

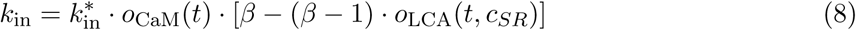

where *k*_in_* is a constant representing the maximum inactivation rate, β > 1 is the increase in inactivation rate when the occupation of o_LCA_ drops due to a decrease in luminal calcium, and o_CaM_ is the occupancy of the Ca^2+^-CaM complex. The dynamics of o_CaM_ is governed by linear binding kinetics of Ca^2+^-CaM of the complex [Ca · CaM] to the binding site:

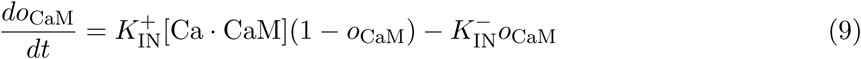

In steady state, this results in a linear dependence of o_CaM_ on the cytosolic calcium concentration, c*_i_*, consistent with the full formulation of inactivation presented by Shannon et al. in [46].

The value of the parameters used in this study can be found in the S1 File. The steady state values of o_CaM_ as a function of c*_i_*, and the steady state values of f*_SR_* = [α + (1 − α) · o_LCA_] and g*_SR_* = [β − (β − 1) · o_LCA_] as a function of free calcium concentration in the SR are presented in Figure 2C and 2D. The global fraction of RyR2 in the open state as a function of free luminal and cytosolic calcium concentration in steady state is presented color-coded in Fig. 2E.

#### Buffering in the SR. Calsequestrin localization

Calcium buffering within the SR is primarily mediated by calsequestrin (CASQ2), which binds calcium rapidly and with high capacity. As shown in Figure 1, CASQ2 is co-localized with the RyR2. To incorporate this effect into our computational model, we make the total buffering capacity of CASQ2 spatially dependent. We define the buffering capacity at each voxel x⃗ as:

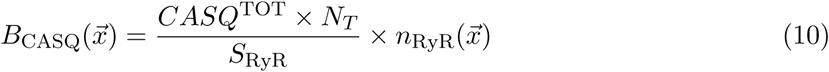

where CASQ^TOT^ is the total amount of CASQ2 in the cell, N*_T_* is the total number of voxels in the myocyte, S_RyR_ is the number of voxels containing a RyR2 cluster, and n_RyR_(x⃗) is an indicator function that takes the value 1 in those volxels where a RyR2 cluster is present, and 0 otherwise.

### Initial calcium load and excitation

The initial state of the system is defined by setting all RyR2 channels and calcium buffers to equilibrium based on the chosen free calcium concentrations in the cytosol and sarcoplasmic reticulum (SR). This fixes the total cellular calcium and the initial occupancy states of the RyR2s, making the total calcium concentration the primary parameter governing the system’s evolution.

Our study focuses on high calcium load conditions conducive to wave propagation. Rather than modeling specific upstream triggers—such as altered NCX equilibrium or increased SR leak—we establish a high total calcium state directly through these initial conditions.

To evaluate the medium’s excitability and its capacity to sustain waves, we apply a localized external stimulus at the cell’s bottom-left boundary (the first three Z-planes). We checked that a minimal external flux or a directed transient opening of the RyR2 leads to the same global behavior. While the model can nucleate waves spontaneously at extreme loads, this “kick” acts as a controlled trigger. It allows us to systematically analyze steady-state propagation dynamics even at calcium levels where spontaneous nucleation is absent, but the medium remains fully capable of supporting a wave. We will differentiate between triggered activity at the initial time and non-triggered activity that happens after this initial trigger.

### Modifications of the physiological model

We performed two in silico experiments to investigate calcium wave dynamics and the functional relevance of CASQ2. First, we compared our physiological model—where CASQ2 co-localizes with RyR2—against a hypothetical scenario in which CASQ2 is uncoupled from RyR2 and distributed uniformly across all voxels. While the total CASQ2 concentration remains identical in both cases, this comparison isolates the effect of high-density local buffering versus a spatially homogeneous distribution.

Second, we explored the impact of NCX conductivity by simulating both down-regulation (knockout) and up-regulation of its function in order to determine how NCX activity affects wave propagation. In our model, down-regulating or knocking out NCX leads to intracellular calcium retention, analogous to hypercalcemic conditions. A complete knockout keeps the initial total cellular calcium constant over time, as the loss of forward-mode extrusion shifts the global equilibrium toward chronically elevated intracellular calcium levels. Conversely, we test whether NCX upregulation enhances calcium extrusion during wave transients, thereby reducing the total calcium load as the wave propagates, or whether it can prevent propagation altogether.

## Results

### The physiological model reproduces experimental calcium wave dynamics

We first tested whether the physiological model reproduces the calcium wave dynamics observed experimentally in isolated mouse ventricular cardiomyocytes. The model incorporates experimentally based RyR2 gating, SERCA activity, cytosolic Ca^2+^ buffering, and the physiological co-localization of CASQ2 with RyR2 clusters. To probe wave propagation, the cell was initialized at a total calcium concentration of C*_T OT_* = 175 µmol/l*_cell_*, corresponding to free cytosolic calcium c*_i_* ≈ 0.2 µM and free sarcoplasmic reticulum calcium c*_SR_* ≈ 1140 µM (Table 1). A localized trigger was then applied at one corner of the cell by injecting calcium into RyR2-containing voxels in the first three Z-planes.

**Table 1.**
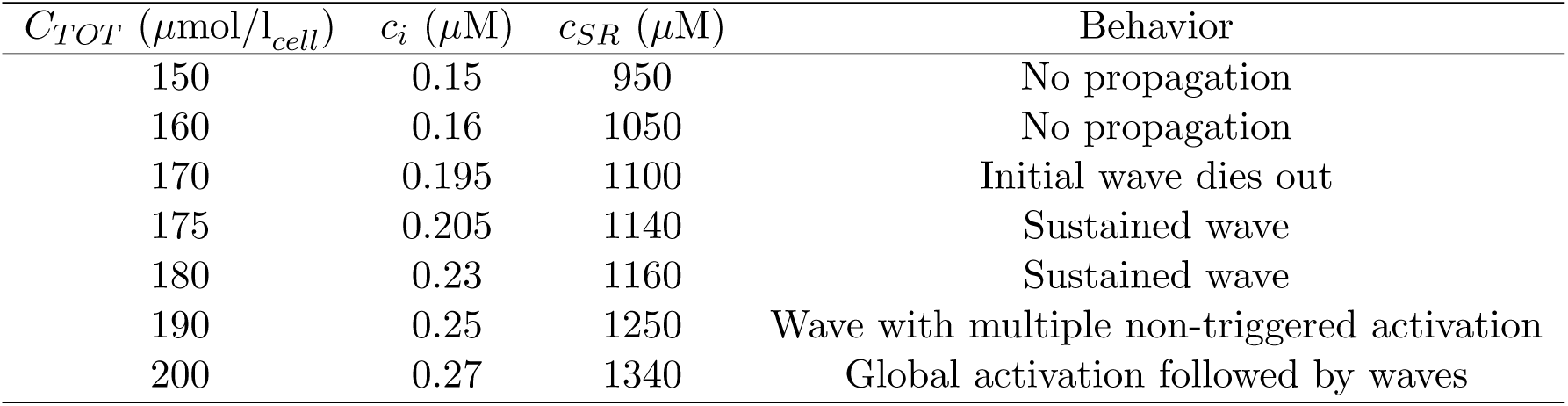
Calcium load regimes and corresponding behaviors in the model.

Under these conditions, the initial activation develops into a coherent wavefront that propagates along the longitudinal axis of the cell by sequentially recruiting successive Z-planes (Figure 3A). The simulated wave reproduces the main macroscopic features observed experimentally: a well-defined front that advances smoothly through the cell without stalling or obvious fragmentation. Sequential frames from the simulation, spanning from 900 ms to 4400 ms after initiation, show the characteristic morphology of the propagating front as it moves through the three-dimensional cellular architecture. The average fluorescence traces ΔF/F*_o_*show an initial rise associated with wave initiation at the left border of the cell, followed by a sustained plateau while the wave traverses the cell, and a subsequent decline once the wave terminates. This agreement indicates that the model captures the large-scale spatiotemporal behavior of the experimental wave.

**Fig 3.**
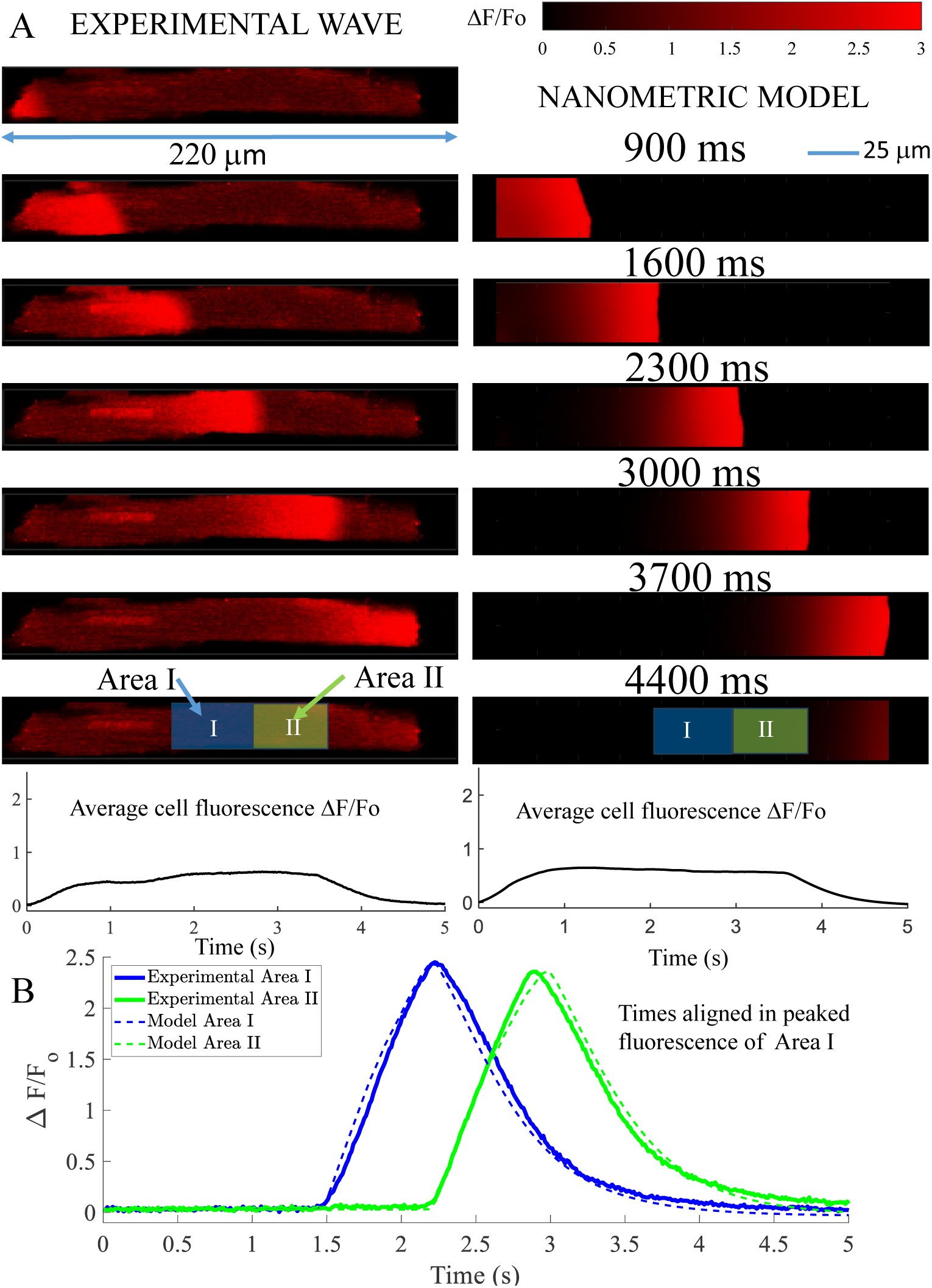
Spatiotemporal comparison of experimental and simulated calcium wave propagation. A) Experimental calcium wave recorded in an isolated mouse ventricular cardiomyocyte compared to the in-silico model. The experimental recording corresponds to the largest cell in our dataset, which exhibited one of the slowest wave propagation velocities. The model simulation was configured with a total cellular calcium load of 175 µmol/l*_cell_*. The wave was initiated via triggered activation at the bottom left corner of the cell grid. Below the images, the average fluorescence of the whole cell exhibits a characteristic plateau during wave propagation. B) Quantitative comparison of fluorescence transients (ΔF/F*_o_*) in two distinct spatial regions (Area I and Area II) along the longitudinal propagation path. Simulated fluorescence was calculated from the concentration of calcium bound to Rhod-2. The time axis for the model traces was shifted so that peak fluorescence in Area I coincides with the experimental peak, facilitating direct comparison of transient kinetics.

The model also reproduces the local kinetics of calcium release and recovery. We analyzed normalized fluorescence traces (ΔF/F*_o_*) computed from the concentration of calcium bound to the fluorescent indicator Rhod-2 in two spatially separated regions along the propagation path (Figure 3B). After aligning the traces at the fluorescence peak in the first region, the experimental and simulated transients show close agreement. The model captures both the rapid upstroke produced by local calcium-induced calcium release and the subsequent decay phase, which is shaped by diffusion, SERCA uptake, and NCX extrusion. The temporal width of the calcium front is also reproduced. Together, these results show that the nanometric model captures both the global progression of the wave and the local kinetics of the fluorescence signal measured experimentally.

### Wave propagation is stable and spans the physiological velocity range

We next examined the macroscopic stability of wave propagation by tracking the wavefront position as a function of time (Figure 4A). In experimental recordings from six isolated cells, the wavefront advances approximately linearly once propagation is established, indicating a nearly constant velocity. The physiological model reproduces this linear progression without evidence of wavefront stalling or collapse. This behavior shows that the spatial arrangement of release sites, luminal buffering, and inter-plane diffusion incorporated in the model is sufficient to support stable propagation over the full length of the cell.

**Fig 4.**
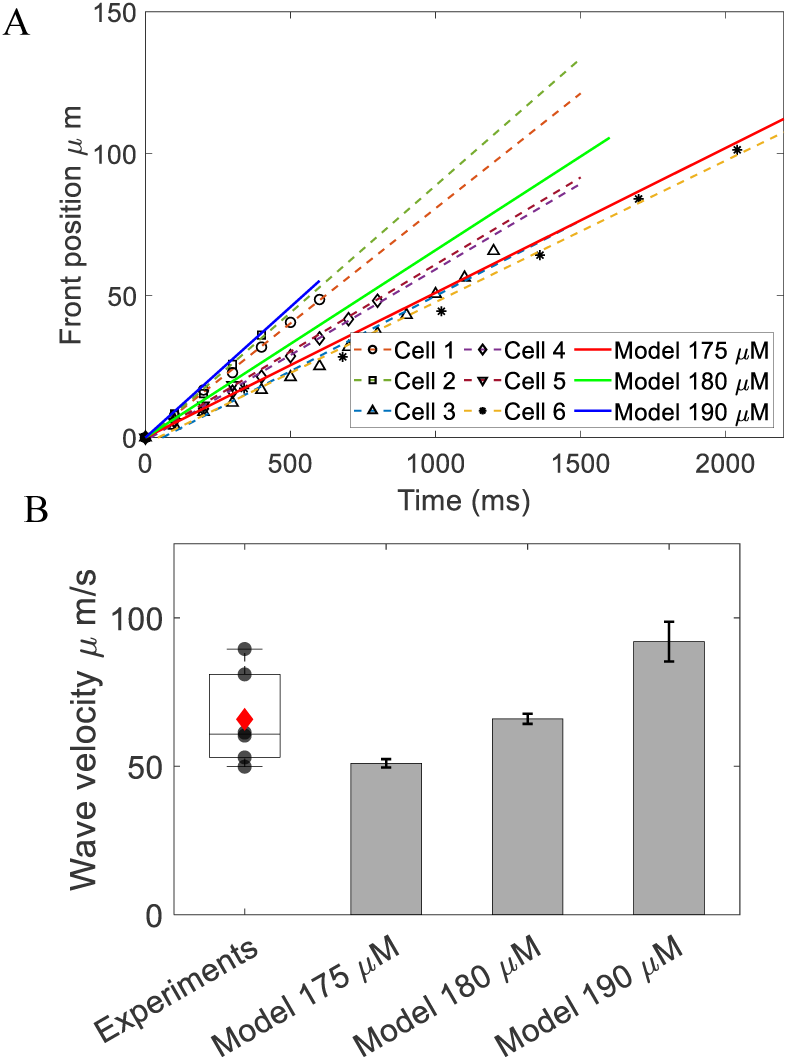
Stability and velocity of calcium wave propagation. A) Time course of wavefront position (µm) for experimental recordings in isolated cardiomyocytes (Cells 1–6) alongside in-silico simulations at three total calcium loads: 175, 180, and 190 µmol/l*_cell_*. The approximately linear progression demonstrates steady propagation without stalling. B) Comparison of wave propagation velocities (µm/s) between experimental measurements and model simulations. By varying total calcium load, the simulated velocities encompass the biological heterogeneity observed in isolated cells.

The model also reproduces the physiological range of propagation velocities observed experimentally (Figure 4B). Experimental wave speeds vary across cells, reflecting biological variability in calcium load and other cellular properties. By varying the total calcium concentration within the wave-supporting regime of the model (C*_T OT_* = 175–190 µmol/l*_cell_*), the simulated velocities span the experimental distribution, falling within the approximate range of 50–100 µm/s. Thus, the model not only supports robust wave propagation, but also captures the dependence of wave speed on calcium loading and the heterogeneity observed across individual cells.

At higher calcium loads approaching 190 µmol/l*_cell_*, both experiments and simulations show that the fastest waves are often interrupted because other regions of the cell activate spontaneously ahead of the primary front. This behavior marks the transition toward global activation at elevated calcium load and is reproduced naturally by the model.

### Calcium load determines the functional regime

The behavior of the physiological model depends systematically on the initial total calcium load (Table 1). Below 170 µmol/l*_cell_*, triggered activations fail to propagate, and the system rapidly settles into a stable balance between SERCA uptake and RyR2 leak. At 170 µmol/l*_cell_*, an initial front forms but dies out after crossing only a few Z-planes. At intermediate loads of 175–180 µmol/l*_cell_*, the system enters an excitable regime in which a localized trigger produces a sustained wave that traverses the full cell. At larger loads, the dynamics change qualitatively. At 190 µmol/l*_cell_*, the triggered wave is accompanied by secondary spontaneous activations at other sites. At 200 µmol/l*_cell_*, the system undergoes global activation, with spontaneous release occurring at multiple locations. These regimes define a clear progression from sub-threshold behavior, to stable propagation, to globally activating dynamics as calcium load increases.

### Uniform CASQ2 distribution abolishes wave propagation

Having established that the physiological model supports robust wave propagation, we next tested whether this behavior depends on CASQ2 co-localization with RyR2. To do this, we constructed a modified model in which the same total amount of CASQ2 is distributed uniformly throughout the sarcoplasmic reticulum rather than concentrated at RyR2-containing voxels. All other components of the model, including total cellular calcium, RyR2 gating kinetics, SERCA activity, and cytosolic buffering, were left unchanged.

Under these conditions, propagation fails (Figure 5). At C*_T OT_* = 175 µmol/l*_cell_*, a calcium load that supports robust propagation in the physiological model, the triggered activation produces a local release that begins to spread but rapidly loses strength. The front becomes pinned and does not advance beyond the first few Z-planes. Even at the higher load of 180 µmol/l*_cell_*, propagation still fails in the uniform CASQ2 model, despite the fact that waves propagate successfully in the physiological configuration.

**Fig 5.**
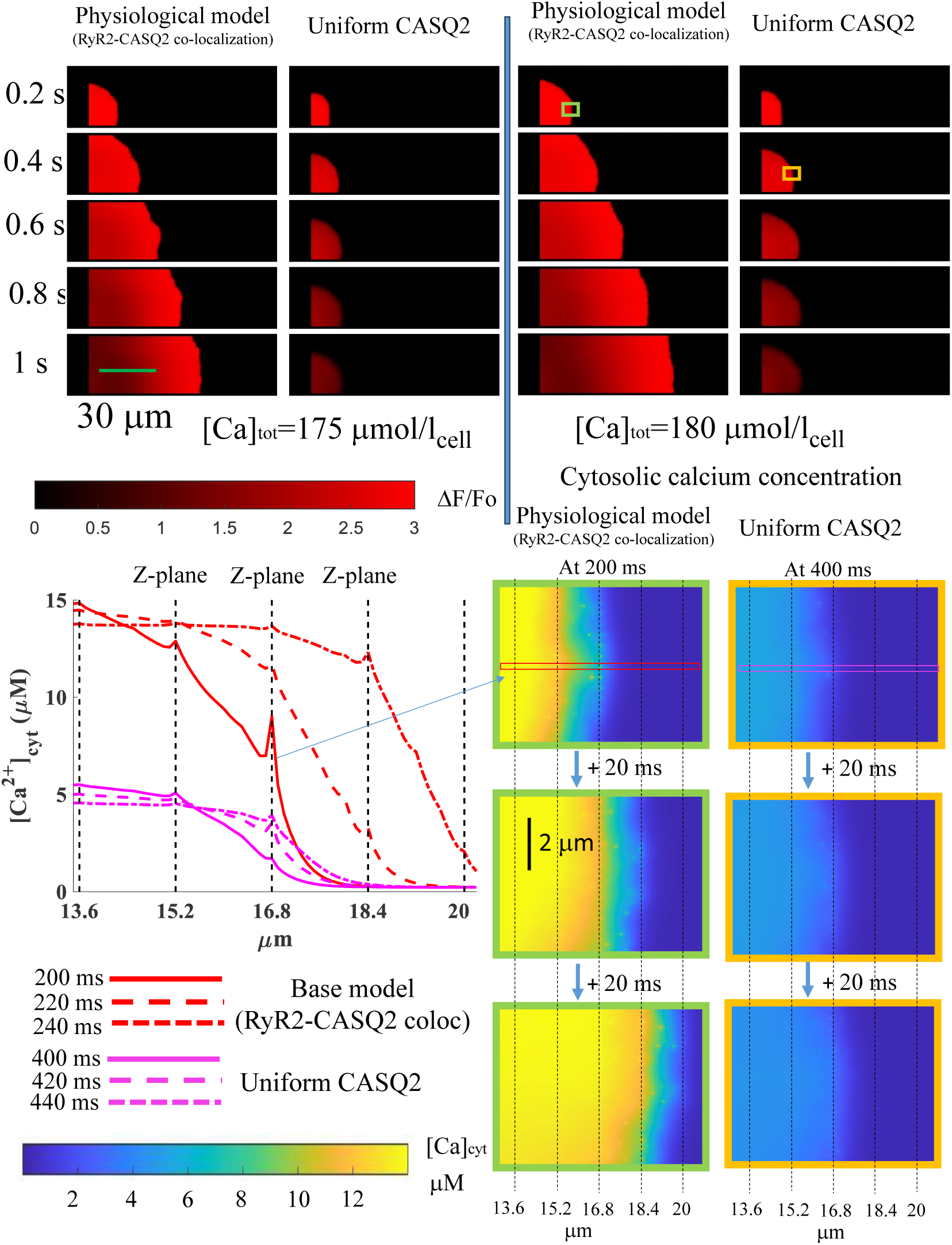
Effect of CASQ2 spatial distribution on calcium wave propagation. (Top) Simulated fluorescence (ΔF/F*_o_*) showing wave propagation across a 90 µm section of the cell over 1 s. Left: physiological model with CASQ2 co-localized at RyR2 clusters. Right: uniform CASQ2 model. Results shown for C*_T OT_* = 175 µmol/l*_cell_* and 180 µmol/l*_cell_*. In the physiological model, the wavefront advances steadily. In the uniform CASQ2 model, the front stalls and fails to propagate. (Bottom) Cytosolic Ca^2+^ concentration profiles at the wavefront, with corresponding line-scan measurements. The physiological model produces cytosolic transients exceeding 10 µM that successfully recruit the next Z-plane. The uniform CASQ2 model produces weaker transients of approximately 5 µM that fail to sustain propagation.

The wavefront analysis reveals why. In the physiological model, the cytosolic calcium concentration at the front rises above 10 µM, well above the half-saturation concentration of 5 µM for RyR2 activation, and the RyR2 opening rate k^+^ remains high in the Z-plane ahead of the front. In the uniform CASQ2 model, the cytosolic transient at the front is markedly weaker, reaching only about 5 µM, and the opening rate ahead of the front is correspondingly reduced. As a result, the calcium signal arriving at the next Z-plane is no longer sufficient to recruit RyR2 robustly, and the front stalls.

The crucial difference is not the total amount of calcium in the sarcoplasmic reticulum, which is the same in both models, but the local release dynamics at the active site. In the physiological model, the concentrated CASQ2 reservoir near RyR2 helps sustain local free SR calcium during release, preserving both the release flux and the luminal support for continued RyR2 opening. In the uniform model, that local buffering capacity is diluted throughout the SR network. The junctional region therefore depletes more rapidly, release weakens earlier, and the cytosolic signal generated at the wavefront is too small to trigger the next Z-plane effectively (Figure 6). Thus, propagation fails not because the cell lacks calcium globally, but because the active release site cannot maintain a strong enough local source.

**Fig 6.**
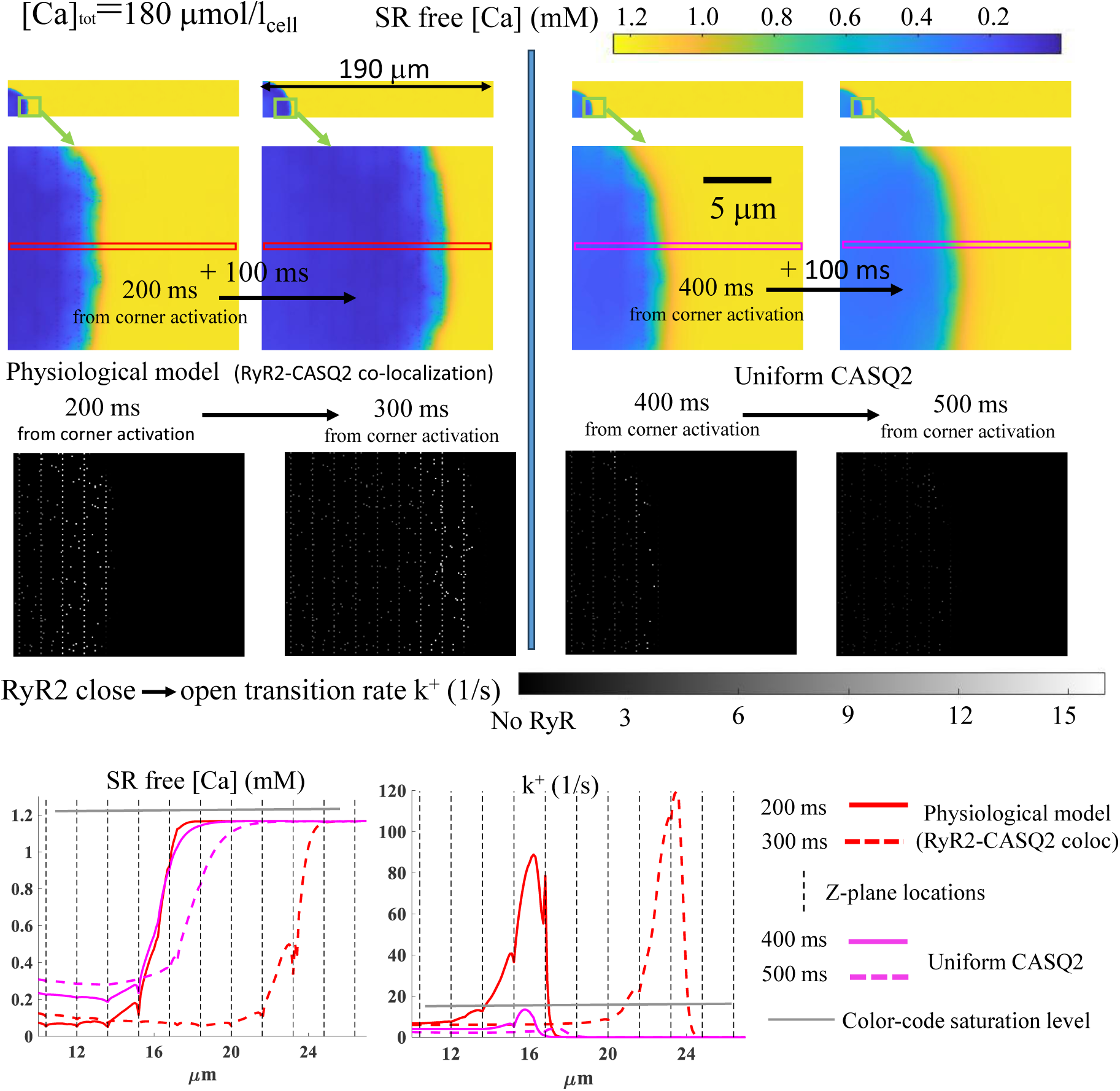
Sarcoplasmic reticulum dynamics at the wavefront. (Top) Spatial maps of free SR Ca^2+^ concentration (c*_SR_* in mM) following triggered activation at C*_T OT_* = 180 µmol/l*_cell_*. Full cell view (190 × 30 µm) with magnified wavefront region (∼15 µm). In the physiological model (left, 200 and 300 ms), the SR depletion front advances longitudinally. Under uniform CASQ2 conditions (right, 400 and 500 ms at equivalent spatial position), the depletion front stalls. (Middle) RyR2 close-to-open transition rates (k^+^, s^−1^) in the wavefront region. The physiological model maintains high transition rates that sustain fire-diffuse-fire propagation. The uniform CASQ2 distribution presents substantially lower rates ahead of the stalled front. (Bottom) Longitudinal profiles of free SR calcium and RyR2 transition rate along the Z-plane. The physiological model (red traces, 200 and 300 ms) shows high k^+^ at the advancing front. Simulations with uniform CASQ2 (blue traces, 400 and 500 ms) show low k^+^ corresponding to the stalled front.

### Role of the sodium-calcium exchanger in wave dynamics and termination

The sodium-calcium exchanger (NCX) is the primary mechanism for calcium extrusion during diastole. Since its activity depends on cytosolic calcium, one might expect wave termination to depend critically on NCX-mediated calcium removal. To test this, we compared the physiological model with two modified cases: a complete NCX knockout and a 16-fold increase in NCX activity. A complete list of the cases studied is provided in S1 File, and all resulting outcomes are available as movies in S2 Github.

Figure 7 compares calcium wave dynamics in the physiological (baseline) model with those under enhanced NCX activity. In the physiological model, the top panel shows representative spatial snapshots for increasing total cellular calcium loads, while the middle panel Figure 7B shows the corresponding evolution of spatially averaged free cytosolic calcium ([Ca]*_cyt_*), total cellular calcium, and whole-cell NCX flux. At low calcium loads (160–170 µmol/l_cell_), sustained propagation fails. At 160 µmol/l_cell_, the initial release cannot propagate along a single Z-plane and rapidly dies out. At 170 µmol/l_cell_, propagation along individual Z-planes becomes possible, but the wave still fails to cross between neighboring Z-planes. NCX extrusion is very low in this case, and total cellular calcium decreases so slowly that it appears nearly constant over the 6-second timescale (See Figure 7B center).

**Fig 7.**
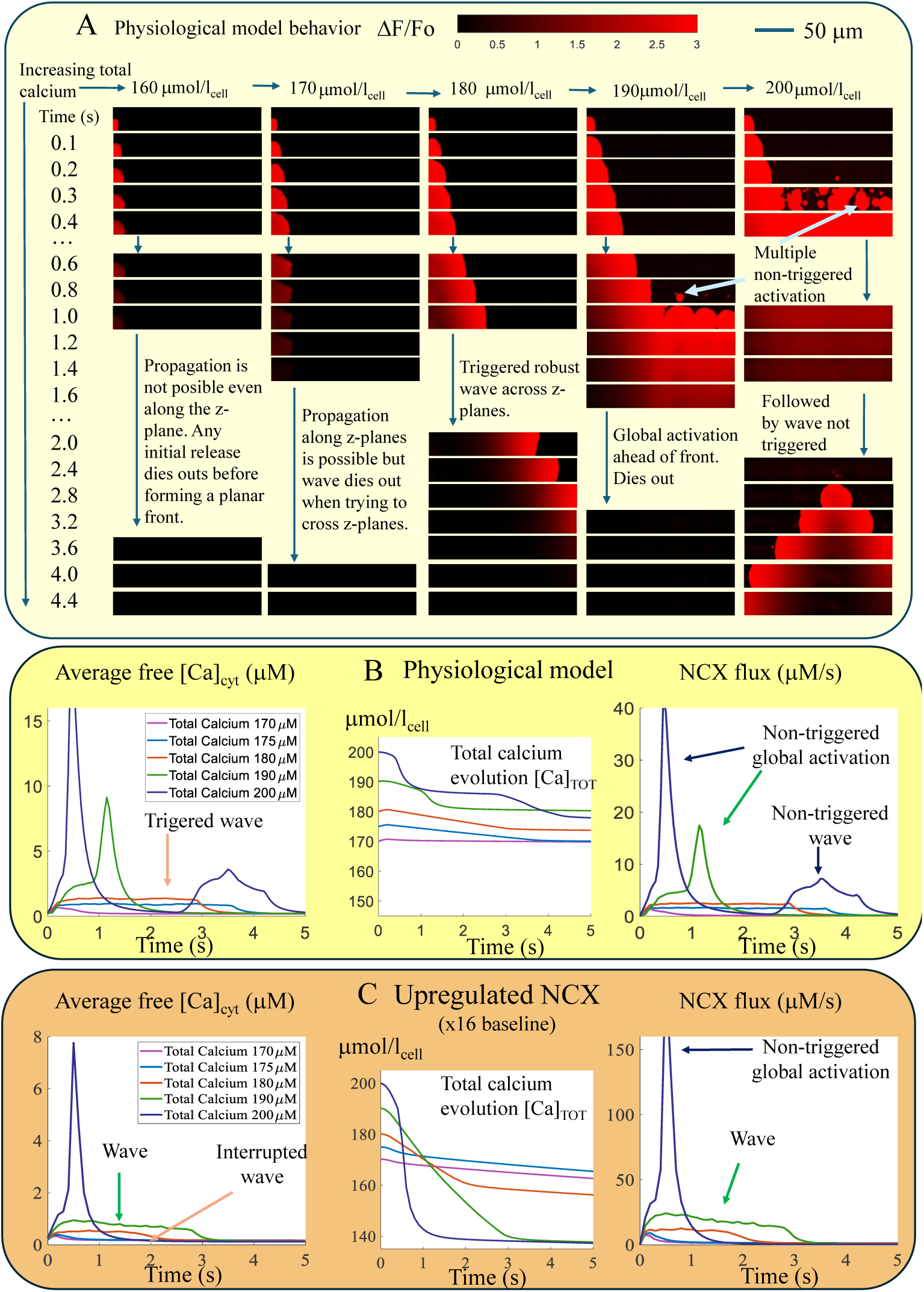
Calcium wave dynamics and the role of NCX-mediated Ca^2+^ extrusion. A) Representative snapshot evolution of cell Ca^2+^-fluorescence at increasing initial Ca^2+^ loads for the physiological model, illustrating the transition from subthreshold response, to stable wave propagation, to multi-site global activation followed as initial [Ca]*_T OT_* increases. B) Time course of spatially averaged free cytosolic calcium ([Ca]*_cyt_*, left), whole-cell NCX flux (center) and total calcium levels of the cell (right) at five initial calcium loads (170, 175, 180, 190, and 200 µmol/l_cell_). At moderate loads, stable waves produce a low plateau in average calcium levels and NCX flux with a slow depletion of the total calcium load. At 190 µmol/l_cell_, spontaneous multi-site activation generates a large transient spike, while at 200 µmol/l_cell_ a global activation is followed by a wave producing a large NCX extrusion and a sharp reduction in total calcium. C) Same time courses for a 16−fold NCX upregulation. Notice that Y-axis s_1_c_6_ales are different than in the physiological model. Enhanced extrusion accelerates recovery and suppresses the transition toward global activation. At 190 µmol/l_cell_, the cell presents now a wave. The reduction of [Ca]*_tot_* over time confirms the enhanced extrusion capacity both with waves and global activation.

At intermediate loads (175–185 µmol/l_cell_), robust waves propagate successfully across successive Z-planes and traverse the full cell. Under these conditions, the average cytosolic calcium and NCX traces exhibit a small plateau during propagation (See Figure 7b left and right), reflecting the fact that only the propagating wavefront is active at any given time. Figure 7B center shows that for 180 µmol/l_cell_, total cellular calcium decreases slowly but progressively because of NCX-mediated extrusion.

At 190 µmol/l_cell_, the initial triggered wave is followed by a non-triggered global activation ahead of the wave front involving most of the cell as seen in Figure 7A. This produces a large calcium transient and rapid calcium extrusion. After this release, the system returns close to the resting state. At 200 µmol/l_cell_, however, the system undergoes immediate non-triggered global activation without formation of an organized propagating wavefront. Under these conditions, cytosolic calcium and NCX flux rise abruptly, while total cellular calcium decreases rapidly due to the large calcium extrusion generated by the globally activated state. This global activation relaxes and a new non-externally triggered wave subsequently emerges. In this case, the wave initiates at the center of the cell. The resulting wave has actually two fronts, one moving to the left and one to the right, producing the characteristic plateau at a higher level than when only one front was present. The calcium extrusion via NCX is smaller than with a global activation but larger than in the case of having one front. After this wave and extrusion, a third wave might appear but eventually, the final state of the system is the resting state with a final total calcium concentration below 180 µmol/l_cell_ (see Figure 7B center)

The complementary simulation with 16-fold NCX upregulation shows a marked change in calcium wave dynamics (Figure 7C left). Enhanced NCX activity increases calcium extrusion during propagating waves, producing larger NCX fluxes and a faster decrease in total cellular calcium during wave propagation (Figure 7C center and right). As a consequence, conditions that generate robust propagating waves in the physiological (baseline) model can produce interrupted waves when NCX activity is strongly increased, indicating that rapid calcium extrusion reduces the calcium available to sustain propagation across successive Z-planes. More importantly, conditions that produce non-triggered global activation events in the physiological model, such as at 190 µmol/l_cell_, are converted into propagating waves when NCX activity is strongly upregulated. However, at the highest calcium loads, the system can still undergo non-triggered global activation. In these cases, the associated NCX flux becomes extremely large, producing a very rapid drop in total cellular calcium. Unlike the physiological model, where global activation can be followed by additional spontaneous activity, the strong calcium loss generated by enhanced NCX activity prevents subsequent reactivation events.

The effects of reduced NCX activity were investigated separately using simulations with complete NCX knockout (Figure 8). In contrast to the significant calcium loss observed with enhanced NCX activity, total cellular calcium remains constant in the absence of NCX-mediated extrusion. Nevertheless, at intermediate calcium loads such as 180 µmol/l_cell_, propagating waves still terminate normally despite the complete absence of NCX-mediated calcium extrusion. The temporal evolution of spatially averaged cytosolic calcium is remarkably similar to that of the physiological (baseline) model, demonstrating that NCX activity is not required for wave termination under these conditions. Instead, intrinsic regulation of the release system, including luminal calcium-dependent RyR2 regulation and calmodulin-mediated inactivation, is sufficient to terminate propagating waves even when total cellular calcium is conserved.

**Fig 8.**
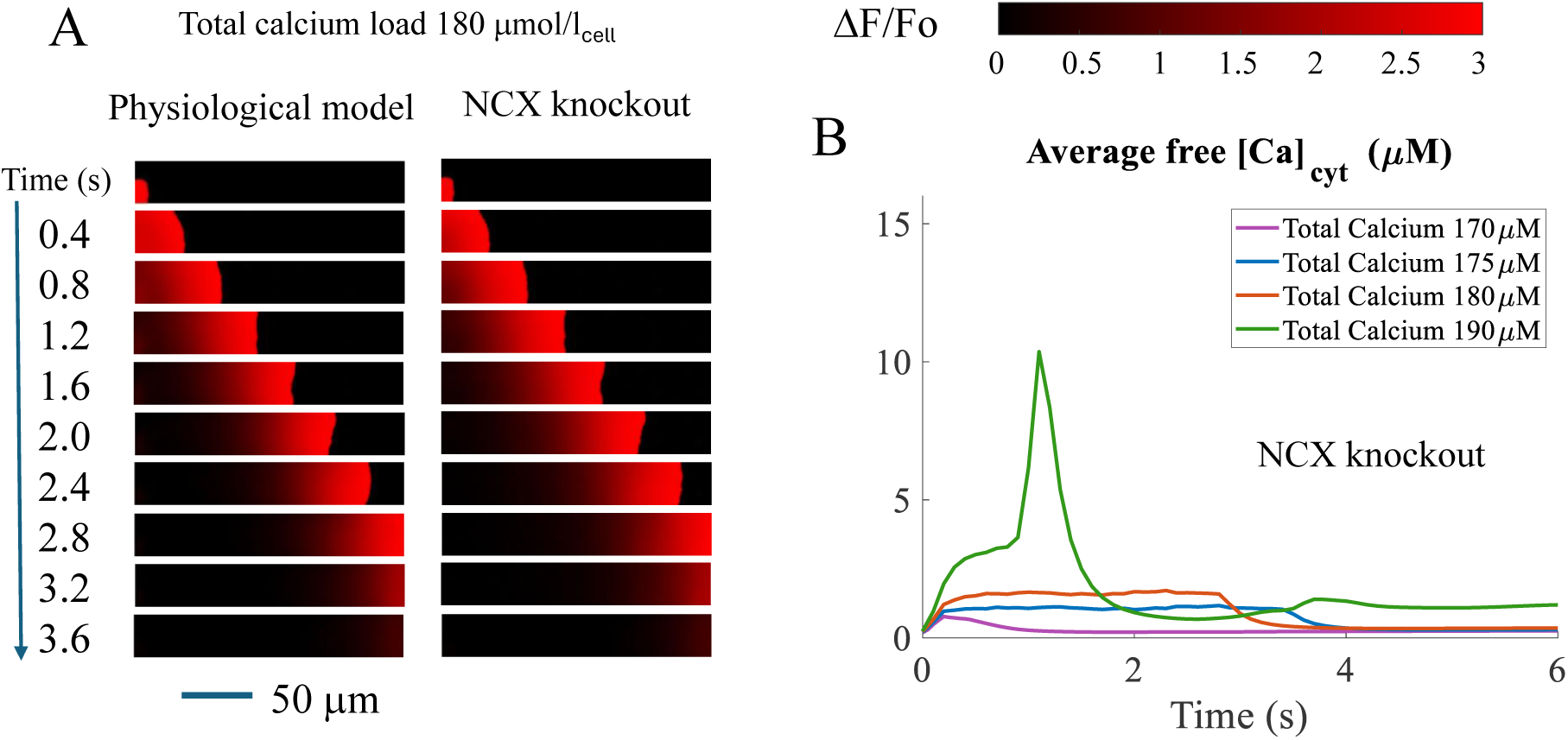
Calcium wave dynamics under NCX knockout. A) Comparative evolution of the physiological model and the model with NCX knockout, for 180 µmol/l_cell_. Wave terminates normally despite the absence of Ca^2+^ extrusion, demonstrating that intrinsic RyR2 regulation is sufficient for wave shutdown. B) Time course of spatially averaged free cytosolic calcium ([Ca]*_cyt_*) for different calcium loads (170, 175, 180, and 190 µmol/l_cell_), with NCX knockout. NCX extrusion is not shown since it is always zero. Total calcium load is also not shown since it is constant. Notice that for 190 µmol/l_cell_ the final state presents a persistent average free calcium around 1 µM and the cell does not return to the resting state.

The importance of NCX becomes clear at higher calcium loads. When total cellular calcium reaches 190 µmol/l_cell_ in the NCX knockout model (See Figure 8B), the system no longer returns to the resting state after the global activation. Instead, the average cytosolic calcium rises and remains at a distinct elevated level where there is a relevant ratio of RyR2 channels partially open and continuous calcium release balanced by SERCA reuptake with a lack of spatial homogeneity. Because there is no calcium extrusion, the cell cannot reduce its total calcium load, and these persistently open RyR2 state remain indefinitely. In contrast, in the physiological (baseline) model after global activation, this state does not appear as NCX-mediated extrusion slowly lowers total cellular calcium, allowing the cell to recover toward the resting state.

Taken together, these results separate two distinct roles in wave control. Intrinsic RyR2 regulation determines whether an individual wave can terminate, whereas NCX controls the long-term calcium inventory of the cell and becomes essential for preventing persistent high-calcium states when total calcium load is excessive. At moderate calcium loads, waves therefore behave as self-limited excitatory events with only modest NCX activation. At higher calcium loads, however, NCX-mediated calcium extrusion becomes critical for eliminating persistent active domains and preventing pathological calcium cycling.

## Discussion

### CASQ2–RyR2 co-localization is a fundamental requirement for propagation

The central result of this study is that the spatial localization of calsequestrin (CASQ2) near RyR2 clusters is required for robust calcium wave propagation in the nanometric ventricular cell model. In the physiological configuration, where CASQ2 is concentrated at release sites, triggered calcium waves propagate across the cell with velocities and overall dynamics that agree with experiment. In contrast, when the same total amount of CASQ2 is redistributed uniformly throughout the sarcoplasmic reticulum while all other components of the model are kept unchanged, the wavefront stalls and fails to cross the cell. This comparison shows that wave propagation depends not just on the total amount of luminal buffer present, but on where that buffer is located relative to the release channels.

This result addresses a fundamental physical difficulty of calcium wave propagation. For a wave to advance, calcium released at one site must produce a signal at the next site that is strong enough to activate RyR2 there. This is difficult because the cytosolic signal is weakened by buffering and geometric dilution as it diffuses away from the source, while the source itself is weakened by local depletion of sarcoplasmic reticulum calcium. Each active site must therefore act as a strong and sufficiently sustained source of calcium despite the fact that release immediately tends to reduce its own effectiveness.

### Localized CASQ2 sustains local source strength at the wavefront

The simulations indicate that CASQ2 co-localization solves this source-strength problem by creating a concentrated luminal calcium reservoir directly at the release site. When CASQ2 is localized to voxels containing RyR2, a large pool of buffered calcium is positioned immediately adjacent to the open channels. As free luminal calcium begins to fall during release, calcium unbinds from CASQ2 and replenishes the local free pool. This delays the fall of free SR calcium, helps maintain the driving force for release, and prolongs the luminal conditions that support RyR2 opening. As a result, the release flux remains large enough for long enough to generate a cytosolic signal that can recruit the next Z-plane.

When CASQ2 is distributed uniformly, this local support is lost. Although total CASQ2 in the cell is unchanged, the amount available near any one RyR2 cluster is much smaller. The junctional SR then depletes more rapidly during release because the nearby buffering capacity is insufficient to sustain local free luminal calcium. Release weakens earlier, luminal support for RyR2 opening falls more rapidly, and the cytosolic signal produced at the wavefront is too small to trigger robust activation in the next Z-plane. Propagation therefore fails not because the cell lacks calcium globally, but because calcium is no longer concentrated where the wavefront needs it most.

This gives a simple mechanistic interpretation of the role of CASQ2 localization. The problem of wave propagation in a buffered medium is fundamentally a problem of local source strength. Each release site must deliver enough calcium, for long enough, to overcome attenuation during transport to the next site. CASQ2–RyR2 co-localization provides a structural solution to this problem by placing a large luminal calcium reservoir exactly where release occurs.

### Implications for wave robustness and for the distinction between sparks and waves

These results have several broader implications. First, they predict that perturbations that weaken the structural coupling between CASQ2 and RyR2 should impair wave propagation even if total sarcoplasmic reticulum calcium content remains relatively normal. Mutations or structural alterations involving CASQ2, triadin, or junctin could therefore affect wave dynamics by changing the local organization of the junctional complex [49, 50], independently of any change in total calcium load. Second, the model predicts that wave velocity and robustness should depend more strongly on the local concentration of CASQ2 at release sites than on average luminal buffering properties alone. Two cells with similar global calcium content could therefore exhibit different wave behavior if they differ in the degree of CASQ2 localization .

The model also clarifies the distinction between calcium sparks and calcium waves. A spark only requires local release in the immediate vicinity of a single RyR2 cluster. A wave requires successful propagation from one release site to another over a much larger distance. Because of this, waves place much greater demands on local source strength than sparks do. A release site may still generate a brief local event even if CASQ2 localization is weakened, but sustained propagation requires a source that remains strong enough to survive cytosolic attenuation and still trigger the next Z-plane.

### NCX is not required for wave termination at moderate loads, but becomes important at high loads

A second important finding concerns the role of the sodium-calcium exchanger (NCX) . The simulations show that calcium extrusion is not required for termination of a propagating wave at moderate calcium loads. In the model, waves initiated at 175–180 µmol/l_cell_ still propagate and return toward baseline even when NCX is completely removed. This shows that wave shutdown is governed primarily by the intrinsic properties of the release system rather than by calcium loss from the cell. As the wave passes, local SR depletion reduces luminal support for RyR2 opening, while calmodulin-mediated inactivation further suppresses release. Together, these mechanisms allow the wave to terminate even though total cellular calcium is not changed.

The role of the NCX becomes more relevant at higher calcium loads. When the total calcium content is sufficiently large, intrinsic shutdown mechanisms are no longer enough to return the cell to a uniform, quiescent state. Under these conditions, the absence of NCX forces the system to remain trapped in a pathological regime characterized by persistent release domains and sustained elevation of cytosolic calcium. In contrast, stronger NCX activity lowers total cellular calcium and suppresses the transition toward global activation. The simulations, therefore, distinguish between two functions: intrinsic RyR2 regulation determines whether an individual wave can terminate, whereas NCX determines whether the cell can progressively unload calcium and avoid a persistent high-calcium state when the overall load becomes too large.

### Self-termination of calcium waves shapes their arrhythmogenic potential

Beyond the mechanistic role of NCX, our results point to the intrinsic ability of calcium waves to self-terminate as a key determinant of their electrical impact. At moderate calcium loads, waves propagate but are rapidly extinguished by local RyR2 regulation, including luminal depletion and calmodulin-dependent inactivation. This self-termination limits the duration and magnitude of calcium release, thereby reducing the amount of calcium that must be extruded via NCX. As a consequence, wave activity in this regime generates only small NCX-mediated inward currents, which are not likely to trigger delayed afterdepolarizations (DADs). For example, in our physiological model, waves produce a flux of around 2-3 µM/s via NCX (see Fig. 7). For typical values of the capacitance-volume ratio in mouse of 7pF/pl [51], this produces an inward current of around 0.04pA/pF. This current is too small to overcome the effect of the outward potassium currents (at around 0.5-1 pA/pF [52, 53]), and thus does not increase the transmembrane voltage beyond the threshold for sodium activation even without taking into account the sodium-potassium pump that further counteracts the effect of NCX. Thus, despite the presence of propagating waves, the system remains electrically stable because the electrogenic effect of calcium extrusion is intrinsically constrained. This interpretation is consistent with experimental observations in healthy versus human atrial fibrillation (AF) myocytes, where all calcium waves occur without leading to triggered activity in healthy cells [54]. In these cells, wave activity remains transient and self-extinguishing, indicating that low NCX engagement during wave termination is not, by itself, pro-arrhythmogenic.

This situation changes fundamentally at higher calcium loads, where the intrinsic ability of waves to self-terminate becomes impaired. Under these conditions, RyR2-mediated release is no longer efficiently shut down, leading to prolonged or recurrent calcium release events. As a result, a larger fraction of released calcium, easily of the order of 40µ M/s (see Fig. 7), must be extruded via NCX, generating stronger inward currents, of the order of 0.4 pA/pF, increasing the likelihood of DADs. The transition to a pathological regime is therefore not simply due to the presence of waves, but to the loss of their self-limiting character, which amplifies their electrogenic impact. In this regime, NCX plays a critical role. It is crucial to terminate the wave and, in order to do so, it has to extrude a significant amount of calcium leading to a sustained and large depolarizing current.

Together, these results point to the fact that the arrhythmogenic potential of calcium waves is determined by the interplay between their intrinsic termination properties and the resulting NCX-mediated current, rather than by wave occurrence alone.

### Limitations

Several limitations of the present study should be noted. First, the model is deterministic and therefore does not capture stochastic RyR2 gating, which could affect the fine structure of the wavefront and the dynamics near the propagation threshold. Second, the comparison between co-localized and uniform CASQ2 represents two limiting cases. Intermediate spatial distributions should be explored in future work to determine how much co-localization is required for reliable propagation. Third, the conclusions are based on a mechanistic computational model, so direct experimental tests of how altered CASQ2 localization changes wave propagation would be an important next step.

Despite these limitations, the main conclusion is clear. In the model, experimentally motivated co-localization of CASQ2 with RyR2 is not a minor structural detail, but a functional feature that allows each release site to act as a sufficiently strong source for the wavefront to advance. This identifies nanoscale luminal organization as an essential component of cardiac calcium signaling.

### Perspectives

Several extensions follow naturally from the present findings. The most immediate concerns the mechanisms that terminate calcium waves. Our results indicate that termination at moderate loads is governed by intrinsic RyR2 regulation, in particular luminal calcium dependence and calmodulin-mediated inactivation. Systematically weakening these regulatory pathways within the model would test directly how the loss of self-termination gives rise to sustained or recurrent release, and how this amplifies the NCX current that drives delayed afterdepolarizations.

A second direction is to examine how global calcium handling modulates these regimes. SERCA activity sets the steady-state calcium load and should therefore influence both the threshold for wave propagation and the stability of self-termination. Exploring a broader range of SERCA activity, along with parameter sets representative of different species or disease states, would clarify how general the mechanisms identified here are, and whether some cellular environments intrinsically favor the transition to persistent calcium release.

A third extension is to couple the model to membrane voltage. The present framework resolves subcellular calcium handling in detail but treats the cell as electrically clamped. Embedding this framework in an electrophysiological model would allow the NCX current generated during wave activity to be translated directly into membrane depolarization, providing a quantitative link between the intracellular regimes identified here and the emergence of triggered activity.

## Conclusion

We have developed a sub-micrometric three-dimensional model of ventricular calcium cycling that identifies CASQ2–RyR2 co-localization as a structural requirement for calcium wave propagation. In the physiological model, where CASQ2 is concentrated at release sites, waves propagate robustly across the cell with dynamics consistent with experimental observations. When CASQ2 is redistributed uniformly while total CASQ2 is kept constant, the wavefront stalls. The mechanism is direct: co-localization provides each release site with a large local luminal calcium reservoir that sustains release long enough to generate a cytosolic signal capable of recruiting the next Z-plane. Without this local support, the source weakens prematurely and propagation fails. These results show that the nanoscale organization of the sarcoplasmic reticulum, specifically the positioning of CASQ2 near RyR2 clusters, is a fundamental determinant of cardiac calcium wave dynamics.

## Supporting Information

### S1 File

Contains complete model equations and parameter tables, numerical integration scheme, additional analysis of wavefront dynamics, and list of supplementary videos of wave propagation.

### S2 Github

https://github.com/ConesaJR/CASQ2_RyR2_waves/. It presents videos of the wave front under different total calcium levels and under different NCX activity. It also presents videos of propagation failure when CASQ2 is uniformly distributed. Finally, the codes are posted there and can be accessed under request.

## Acknowledgments

We thank the participants of the KITP program ’ Multi-Scale Physics of Normal and Diseased Heart: from Ion Channels to Whole Organ’ for valuable discussions. This research was supported in part by grant NSF PHY-2309135 and the Gordon and Betty Moore Foundation Grant No. 2919.02 to the Kavli Institute for Theoretical Physics (KITP). This work was supported by grant from Spanish Ministerio de Ciencia, Innovacíon y Universidades (PID2023-152610OB-C21/22 funded by MCIN/AEI/10.13039/501100011033) and grant 2021 SGR 00582 funded by Agència de Gestío d’Ajuts Universitaris i de Recerca (AGAUR).

